# How Should Eukaryotic Chemotaxis be Measured?

**DOI:** 10.1101/2023.08.25.554886

**Authors:** Luke Tweedy, Peter A. Thomason, Robert H. Insall

## Abstract

Chemotaxis and directed cell migration are fundamentally important to eukaryotic biology. To understand the mechanisms that drive such complex processes, informative and robust measurements are essential, but the field does not always agree what these should be. Here we identify the most dependable measures of chemotactic steering and the underlying cell migration, and provide tools to test them. Some widely-used metrics can end up misleading, in particular “cos θ” for directional accuracy. We suggest that chemotactic efficiency should be used as a primary metric. Mean squared displacement and directional autocorrelation can be used to unpick different models of random and directed cell migration. Transition matrices are another useful tool for understanding migration mechanisms and avoiding artefacts, and provide a graphical illustration of how well cells maintain each direction. Unexpectedly, the choice of start and end points of tracks strongly affects the measurements and can seriously bias the measured results. This is particularly clear when cells are not homogeneously distributed at the start of an assay. To support straightforward adoption of these methods, we provide a suite of tools as a plugin for the open-source ImageJ program, and describe how they can be used to understand complex scenarios like self-generated chemotactic gradients.

## Introduction

Many cells need to move to fulfil their biological functions. During embryonic development, many cell types begin to differentiate far from their intended adult site, and must therefore migrate for long distances (Aalto et al., 2021, Aman and Piotrowski, 2010, Starz-Gaiano et al., 2001). Mutations affecting cell migration cause many congenital problems including reduced fertility (Knaut et al., 2003), pigmentation defects (Li et al., 2011), and life-threatening conditions such as Hirschsprung’s disease (McCallion and Chakravarti, 2001). Directed migration is required in to maintain adult health, for example in wound healing (Schneider et al., 2010) and response to pathogens (Xu et al., 2021). It is also a defining feature of cancer metastasis (Muinonen-Martin et al., 2014). As such, a proper understanding of why and how cells move is crucial to tackling cancers and chronic inflammatory diseases. Chief among the many mechanisms cited for steering cells is chemotaxis (the directed migration of cells in response to a chemical gradient; Buracco et al., 2019, Devreotes and Zigmond, 1988). Given both its importance to biology, and the complexity of the processes it is involved in, it is vital that we properly understand and interpret experimental chemotaxis data. There are many methods for quantifying chemotaxis; here we will discuss the merits of several prominent examples, as well as their limitations in the face of real biology and practical, experimental circumstances. We comment on when ensemble statistics may not be appropriate for representing results. Finally, we offer a comprehensive package in the form of an ImageJ (Schneider et al., 2012) plug-in, which will calculate alternative statistics and perform several informative analyses, as well as performing fast automatic tracking and providing field standards such as Cos(8) and cell speed.

### Direct View Chambers

Directed cell migration, particularly chemotaxis, can be measured using a variety of experimental techniques. The most effective method entails observing movement in a direct view chamber. Examples of such devices include the Zigmond (Zigmond, 1977), Dunn (Zicha et al., 1991), and Insall (Muinonen-Martin et al., 2010) chambers, all of which allow a time lapse recording of cell migration in a thin viewing area between two large reservoirs. Typically, one reservoir is filled with untreated medium, and the other medium with added attractant, creating a simple gradient with a defined direction. Such assays provide rich information on the cells’ behaviour, but they are relatively slow to set up, create relatively large volumes of data, and do not allow high-throughput screens.

Chemotaxis is also assayed using end-point assays, in which cells are examined after a fixed interval. The ‘two-spot’ assay, for example, examines the change in the distribution of cells in a tiny liquid droplet following the addition of a similar attractant droplet on one side (Konijn, 1965). The two-spot assay is also difficult to set up and is highly sensitive to humidity, which can cause the droplets to swell and merge or to dry out entirely. Another endpoint approach is the ‘Boyden’ or transwell assay (Boyden, 1962), in which cells are settled on a membrane across which a putative attractant can be sensed. Chemotaxis is then scored by how many cells cross the membrane towards medium with an attractant. This is easy to multiplex, but the information is restricted to a single number for each experiment; a variety of biological processes can cause false positives, for example if the putative attractant alters adhesion or receptor expression. Direct-view chambers are therefore strongly preferred for mechanistic studies.

At present, statistical reporting in chemotaxis often uses Cos(8), the cosine of the angle between a key direction (usually an experimentally imposed attractant gradient) and the overall direction of the cell. A key advantage of Cos(8) is its ease of interpretation-it has a natural range of -1 to 1, with these values meaning respectively that the cell moves directly down or up the gradient, and zero meaning there is no sign of directional bias along the gradient axis. However, there are many disadvantages to this statistic. Two fundamentally different methods go by the same name. In the first, θ is only measured between the start and end of each cell’s track. In the second, cells are imaged at fixed intervals; the θ is measured at the start and end of each interval, then averaged for each cell. Both interpretations are subject to (different) errors at the level of the individual cell, described below, which will add noise. Finally, the resulting value can vary greatly depending on the sampling or frame rate.

## Results

### Chemotactic efficiency index is a better measure of chemotaxis than Cos(8)

Both ways of measuring Cos(8) – the mean Cos(8) across all frames, and the Cos(8) of the line between the beginning and end points of the track – can give flawed measurements of chemotaxis, as shown in Figure 1.

**Fig. 1:**
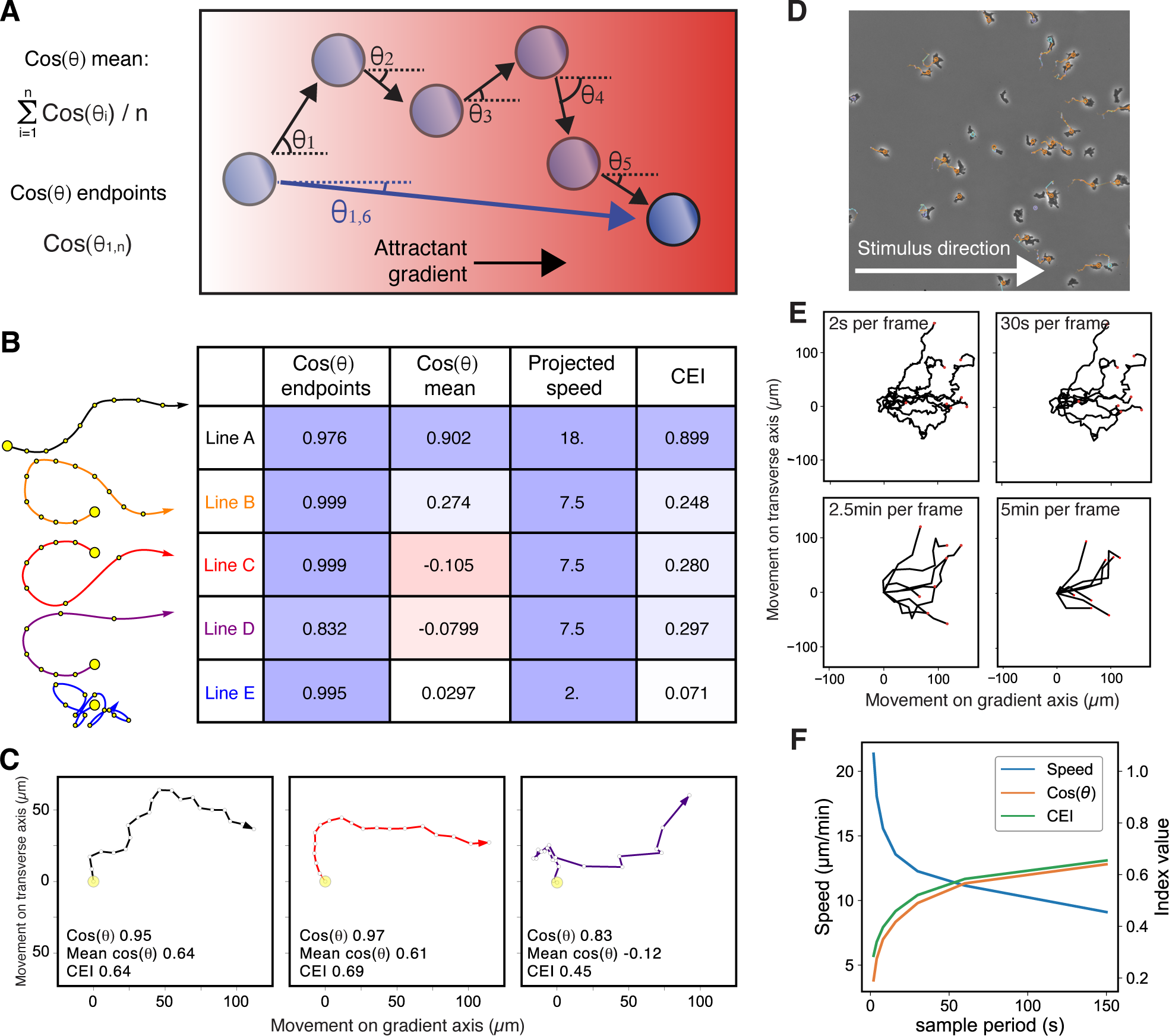
Chemotaxis Efficiency Index is more effective than Cos(8) as a measure of steering. **(A)** Diagram showing two different ways in which Cos(8) is calculated. **(B)** Schematic showing different types of path. Sampling points are shown in yellow. The first path maintains its direction well. The second, third and fourth steer toward the correct direction (rightward), with the third and fourth also increasing speed as they reorient. Example 5 does not follow the gradient, moving little and noisily. **(C)** Computational models of different chemotaxis parameters explain the variance between CEI and Cos(8). Left: moderate orientation, moderate persistence, constant speed: CEI and Cos(8) are the same. Middle: moderate orientation, strong persistence, moderate chemokinesis: CEI is higher than Cos(8) because cell migrates more rapidly when it has time to orient. Right: no directional orientation, weak persistence, strong direction dependence on speed: CEI is positive because migration up-gradient is much faster than down-gradient. Cos(8) is negative because cell spends more time moving down-gradient; but because its speed is slower, downgradient migration contributes less to net distance. **(D)** Example of *D. discoideum* cells being tracked in a 1-well under agarose assay. This movie is deliberately oversampled, with imaging using a 20x phase contrast objective and frames every 2s. **(E)** A collection of the tracks from (C) are shown, sampled every 2s, 30s, 2.5 min and 5min. Note that the tracks sampled every 2s are very tortuous, where the tracks sampled every 2.5min or 5min are very straight and simplified. **(F)** Values of cell speed (blue), Cos(θ) (amber) and chemotactic efficiency index (CEI -green) are shown as a function of sampling period. We encourage a sampling period in which cells move 1-2 cell lengths as this avoids both oversampling noise (shape change and technical centroid position noise) and undersampling noise (removing significant parts of the cell’s movement). In this case, this would be between 30 and 60s.

The principal aim is to quantify how accurately a cell can migrate in the direction of the stimulus (in chemotaxis, this is towards the highest chemoattractant concentration). Our measurements therefore aim to analyse the relationship between this direction and the detailed movement of the cell. Unfortunately, migration is complex, and it is not always clear what constitutes the best relationship. Consider the five example tracks in Fig. 1A. Track E (blue) has no relationship with the gradient direction and is unambiguously poor. Track A (black) has a consistently good alignment with the correct direction of motion. Tracks B, C and D (respectively, orange, red and purple) are all similar - the initial direction is wrong, but the cell corrects it and ends up aligning with the gradient, but with purple maintaining alignment better once the cell reorients. One question is therefore whether average alignment or a tendency toward correct alignment should feature more strongly in the overall assessment. As a reported statistic, Cos(8) does not describe how efficiently errors are corrected - the assumption underlying most chemotaxis assays is that cells are effectively in an equilibrium state, in which cells have already had the time they need to reorient to the externally imposed signal, and that we are now measuring the limits of their ability to read and maintain this chosen direction. This presumes that the external stimulus is constant in direction, strength and degree of noise, a presumption which is usually biologically incorrect – most physiologically relevant cases of chemotaxis happen in complex environments, and many chemotactic systems (for example self-generated gradients; Tweedy and Insall, 2020) are fundamentally out of equilibrium.

However, if we do take general alignment to be the key indicator of successful chemotaxis, we must seek a tracking statistic that ranks the tracks in Figure 1A as A > B, C, D > E. Endpoint Cos(8) does not achieve this ranking, getting instead B, C > E > A > D. In particular, it reports track E as being nearly perfect chemotaxis, despite most of the cell’s decisions being essentially random. The more data-rich mean Cos(8) does rectify that track A is a clear leader. However, it gives both tracks C and D a negative value. This implies that cells are migrating away from the chemoattractant source, even though in reality they are constantly improving their direction and end up travelling in the correct direction (Fig. 1B). This inaccuracy is caused by the cells speeding up when heading up the gradient, which is not only biologically reasonable, but true for many types of chemotactic cell.

An alternative statistic in current use is the chemotactic efficiency index (CEI; Singh et al., 2022). This is sometimes referred to as simply the chemotactic index, but the term is ambiguous – as some users refer to Cos(8) as chemotactic index, we avoid the term here. The CEI is the component of the distance travelled up the gradient divided by the total distance travelled. It shares the (-1,1) range of Cos(8), as well as the physical interpretation of those values, but crucially accounts for the cases that Cos(8) has difficulty with. The CEI ranking of the tracks is, as desired, A>B, C,D>E (Fig. 1B).

The difference in accuracy between mean Cos(8) and the CEI depends to a large extent on the mechanisms underlying chemotaxis. Figure 1C shows three examples. On the left, cells move at a constant speed; the direction of the chemoattractant gradient determines only the direction of the cells as they move. Under these conditions, mean Cos(8) and CEI are essentially identical. However, this is an unrealistic model of nearly all crawling chemotaxis. The centre shows an attractant that causes chemokinesis – that is, if the cells move faster when the concentration is higher – so cells tend to travel further when they get nearer the attractant source; chemotaxis becomes more efficient. The CEI accurately reflects this change, whereas mean Cos(8) ignores it. The majority of chemoattractants also act as chemokines; in immune cells the predominant family (CCLs and CXCLs) were originally named chemokines.

The clearest difference, however, occurs when cells travel faster if they are steered in the correct direction(Fig. 1C, right). When this mechanism is in play, the CEI and mean Cos(8) may even have opposite signs; if cells spend 70% of their time oriented inaccurately, but move three times as fast when migrating up-gradient, they will reliably migrate towards the source despite a negative mean Cos(8). Thus if cells change their speed according to the gradient, mean Cos(8) becomes a seriously flawed measure of steering. In our experience (for example Tweedy et al., 2013) this is very often the case for chemotactic cells.

In some circumstances, a reasonable alternative to CEI is the projected speed (this is the velocity component at any time in the direction of the gradient). This statistic maintains the same score order as CEI, but it loses the natural -1 to 1 range and is not dimensionless, making comparisons across cell types difficult. Of course, comparisons across cell types must be made with great caution – a CEI of 0.3 would be very weak for neutrophils or *Dictyostelium*, but remarkable for melanoma or breast cancer cells. Nevertheless, projected speed downplays tiny, noisy movements, and is a good choice where experimental conditions are complex, for example if changes in attractant have chemokinetic as well as chemotactic effects.

These tracking statistics are all dependent on the rate at which the track is sampled (in practical terms, the frame rate). Errors can be introduced by poor choice of frame rate and by noise in the measurement of cells’ positions. Fig 1D shows cells being tracked at a higher frame rate and magnification than is useful for determining such statistics. When we examine the tracks from this assay at different sampling rates (Fig 1E), we can see that high frame rates emphasise cell shape changes over actual migration, as well as exacerbating technical noise (e.g. jitter in the detected “centre” of the cell). In contrast, low frame rates (where cells move more than 2 cell lengths between frames) may miss substantial parts of the true track. While we sub-sample the automatic track, a true low framerate movie could also be difficult to analyse, with cells often confused with one another. In general, high frame rates tend to inflate apparent speed and decrease alignment indices (such as CEI), with low frame rates showing the converse (Fig. 1F). This has also been observed as a general bias in published reports of cell speed and chemotactic index (Frattolin et al., 2021). In our experience, an ideal frame rate allows cells to move ½ - 2 cell lengths per frame. Where a high frame-rate is required for other experimental reasons (for example, when co-assaying direction of motion with cell shape or a cytoskeletal probe), an appropriate subsample of the frames should be used for calculating CEI.

We suggest that CEI should be a first choice metric as a basic measure of the accuracy of chemotaxis. Projected speed may be favourable where comparisons use the same cell type, and chemokinesis or random migration are prominent in at least one condition. Cos(8) is a much less useful measure. As traditionally used, it is ambiguous whether endpoint or mean Cos(8) is being reported; endpoint Cos(8) is a crude and uninformative measure; and mean Cos(8) is often flawed under physiological conditions.

### Model fitting yields informative migration metrics with a strong foundation in theory

One excellent solution to the errors described above is to fit a model behaviour to the measured data, an approach favoured by mathematical biologists (Gunawardena, 2014) as - unlike our previous summary statistics-the parameters generated are usually robust to changes in sampling rate. Model fitting also provides a theoretical framework for understanding cell behaviour (Metzner et al., 2015, Taylor et al., 2013, Wu et al., 2015), and thus can offer some mechanistic insight. In particular, tracks are often characterised by their persistence and bias, as random walks with these features have well-characterised behaviours and properties (Wu et al., 2015). The presence or absence of persistence and bias combine to give four types of walk, which need to be distinguished to understand cell migration parameters.

At its simplest, a cell may move with a random walk (RW)-that is, frame to frame we expect a cell to move at a consistent speed in a randomly selected direction with no particular weighting. In a random walk the cell has no net direction (Fig 2A, B). The speed may vary from frame to frame, but keeps close to the mean as it does not depend on any external factor. If, on top of this random movement, a cell favours a direction (if it is chemotaxing for example), the result is a biased random walk (BRW, Fig. 2C, D).

**Fig. 2:**
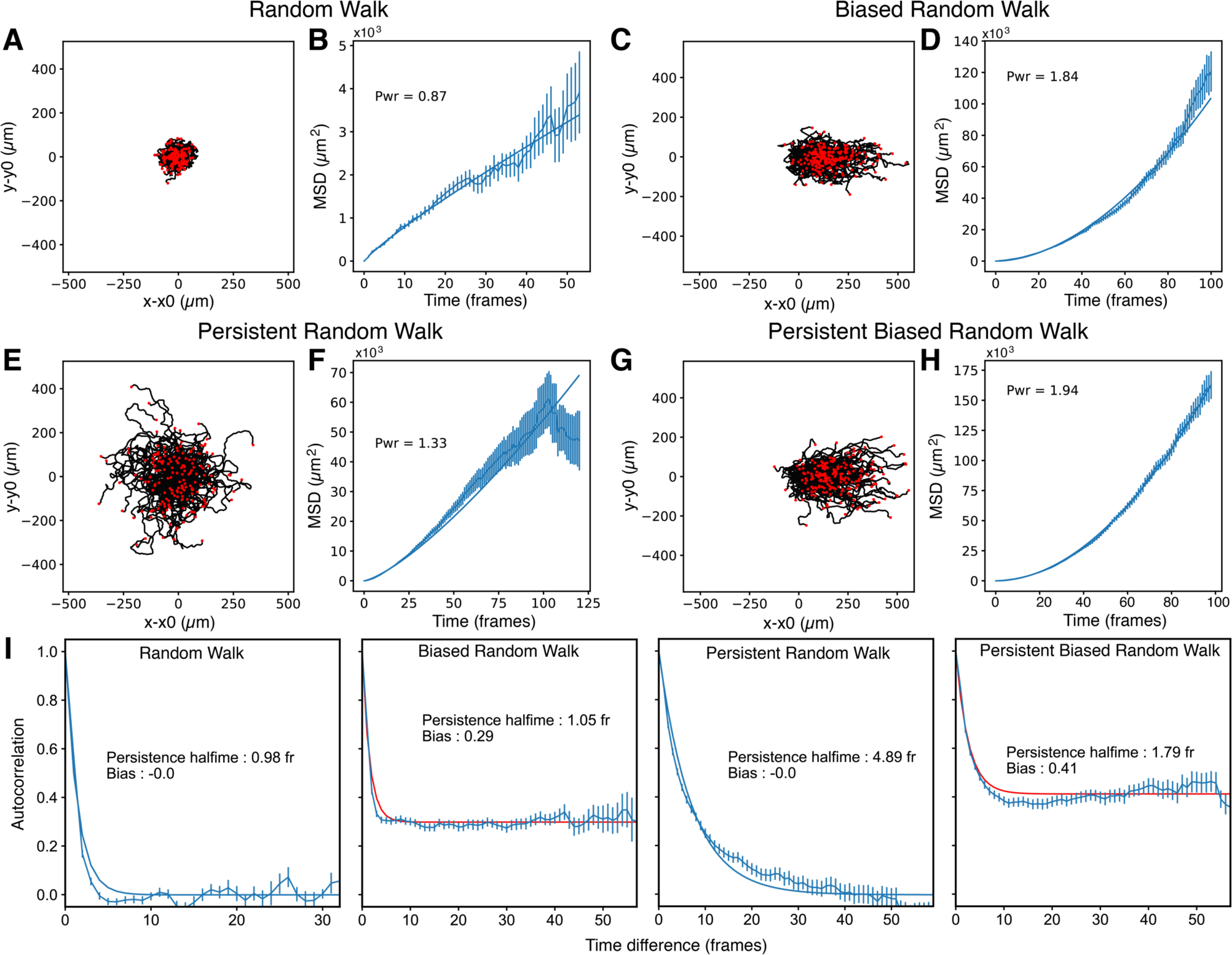
Model fitting is a robust alternative to directional measurement. In this figure, we have generated several behavioural extrema by simulation to demonstrate the uses of model fitting. **(A)** A spider plot of a random walk (RW). The spider plots shows the tracked movement of each cell relative to its starting location. **(B)** Mean squared displacement (MSD) of the random walk. In a random walk, the cell’s direction of motion at any given time bears no relation to its direction at any other. As such, its movement is diffusive (moving outward with the square root of time), and its MSD is approximately linear. We have fitted an MSD (*y* = *Ax^b^*) to the ensemble (red line). Here movement is sub-diffusive (the exponent *b* of perfectly diffusive walk has a value of 1), as more mobile tracks are lost from the field of view. This is a realistic factor in wet lab experiments that we have maintained in the simulated example. **(C, D)** Spider (C) and MSD (D) for a biased random walk (BRW). The directional element raises the exponent nearer to 2 (2 would represent motion in a straight line at a constant speed). **(E, F)** The same for a persistent random walk (PRW). As cells tend to persist in one direction they move larger distances overall, and have an exponent value near two early on (during their persistence time), after which the exponent tends toward 1. With sampling periods longer than the persistence time, a PRW is a RW with a higher diffusivity, which may be misinterpreted as a higher speed. Persistence can be determined by fitting autocorrelations across increasing time periods and seeing if the value remains approximately the same, or if it decays as time goes on. **(G, H)** The same for a persistent biased random walk (PBRW). Like the BRW, the PBRW has a fit exponent close to 2 as the cells move consistently away from their starting location. If the bias is strong it can make it difficult to determine persistence using the MSD, as the exponent does not decay over time. **(I)** Autocorrelation of the four random walks. Autocorrelation, here, means the alignment of the cell direction with its future self, for a range of time intervals. Bias introduces a minimum alignment to which the track converges over longer time periods (contrast the RW with the BRW and the PRW with the PBRW). Persistence can also be determined, as it elongates the time to converge. Here, we fit a time to half-maximal alignment in each case. In the RW and the BRW, persistence half time is ∼0.5-that is half a frame! There is no correlation after one frame. The PRW has a half time of 4 frames, correctly identifying the persistence factor in the simulation of 0.85 (0.85^4^ = 0.522). However, persistence and bias interact-the PBRW has both a higher bias than the BRW and a lower persistence than the PRW because persistence keeps cells aligned with the bias direction, raising the apparent minimum and reducing the time taken to converge on it. The fit nonetheless does report a persistence time higher than one frame.

The two walks are easily distinguished by assessing the net distance cells travel as a function of time. This is because cells using a random walk only increase their net migration distance in proportion to the square root, because they often go back on themselves, so after travelling any given distance, getting twice as far takes four times as long. In comparison, if the random walk is biased, the distance travelled in the direction of the bias increases linearly with time - in twice the time they get twice as far. Practically, this means random migration can move cells a significant distance over short timeframes, but it gets less and less efficient over longer times. Random walks are extremely inefficient at covering a substantive distance; for this, some directed migration is essential.

The standard way to measure this difference is the mean squared displacement (MSD). As the displacement of a perfectly steered cell ensemble goes up linearly over time, its MSD increases linearly in proportion to the square of time, whereas for a group of random walkers MSD is proportional to time. One can therefore fit a function y =a t^x^ to the MSD for an experiment to determine how much movement is randomness (where x = 1) and how much is bias (x = 2). Of course, these walks are not discrete classes, but describe a range of behaviours, so the fits will be approximate.

More realistically, cells might favour their current direction of motion over some time-frame. A lamellipod or pseudopod, one started, continues to direct the cell in one direction for its whole lifetime, and the directions of one dominant pseudopod and its daughter are known to be correlated (Andrew and Insall, 2007). Similarly, the patterns of cytoskeletal activators like active Rac1 are slow to change. If these factors influence a cell over a longer time-frame than the sampling rate, cell direction will be correlated from one frame to the next. This leads to a persistent random walk (PRW, Fig 2E). Persistence can affect the fitting of an experiment if not accounted for (Fig. 2F) as the continued movement of the cell in one direction over a characteristic ‘persistence time’ can resemble directional bias in a plot of MSD. Beyond the persistence time this effect vanishes, with the linear MSD of a random walk establishing itself (see (Gorelik and Gautreau, 2014) for a discussion of this).

A persistent biased random walk (PBRW) combines persistent and biased migration. The two effects can work with or against one another depending on the direction of the cell; persistence causes misalignments in the cell’s direction of motion to last longer, but also allows for a smaller bias signal to keep cells well aligned with the attractant gradient. Persistence can, in fact, be seen as a mechanism for filtering noise in gradient directional sensing (Aquino et al., 2014). As a PBRW is biased walk, the measured MSD increases with the square of time (Fig 2 G, H).

A different way to separate persistence and bias uses directional autocorrelation (Gorelik and Gautreau, 2014). Persistence by definition affects direction over the short term, but not long term. Bias, however, influences direction consistently, over both short and long terms. It is therefore possible to separate two parameters by correlating the current and past direction of a cell’s path. Persistence, over which a cell’s current and past alignment correlate, decays with a characteristic half-time. Bias, which is constant, may therefore be captured in the degree of correlation left over after the persistence time. Fig. 2I shows the autocorrelation fits for our four canonical random walks, and again provides a way of expressing the proportions of cell movement that are random and due to bias. In order to separate these parameters reliably, a group of cells must A) be tracked for longer than the persistence time, and B) include a sufficient number of randomly aligned cells in order that persistence and bias can lead some cells in different directions.

To illustrate the application of these fits to real data, we examined two experimental cases that fit very well to two canonical types of walk. We examined starved *Dictyostelium discoideum* cells responding to a directional stimulus (10µM cAMP, one well under agarose assay, Fig. 3A) or a uniform, non-degradable stimulus (1µM Sp-cAMPS, uniform cell distribution under an agarose slab, Fig. 3B). MSD (Fig. 3C) and autocorrelations (Fig. 3D), both demonstrate that the homogeneous stimulus is best described by a PRW, and a directional stimulus by a PBRW. Both measurements are reasonably robust to changes in frame rate (Fig. 3E).

**Fig. 3:**
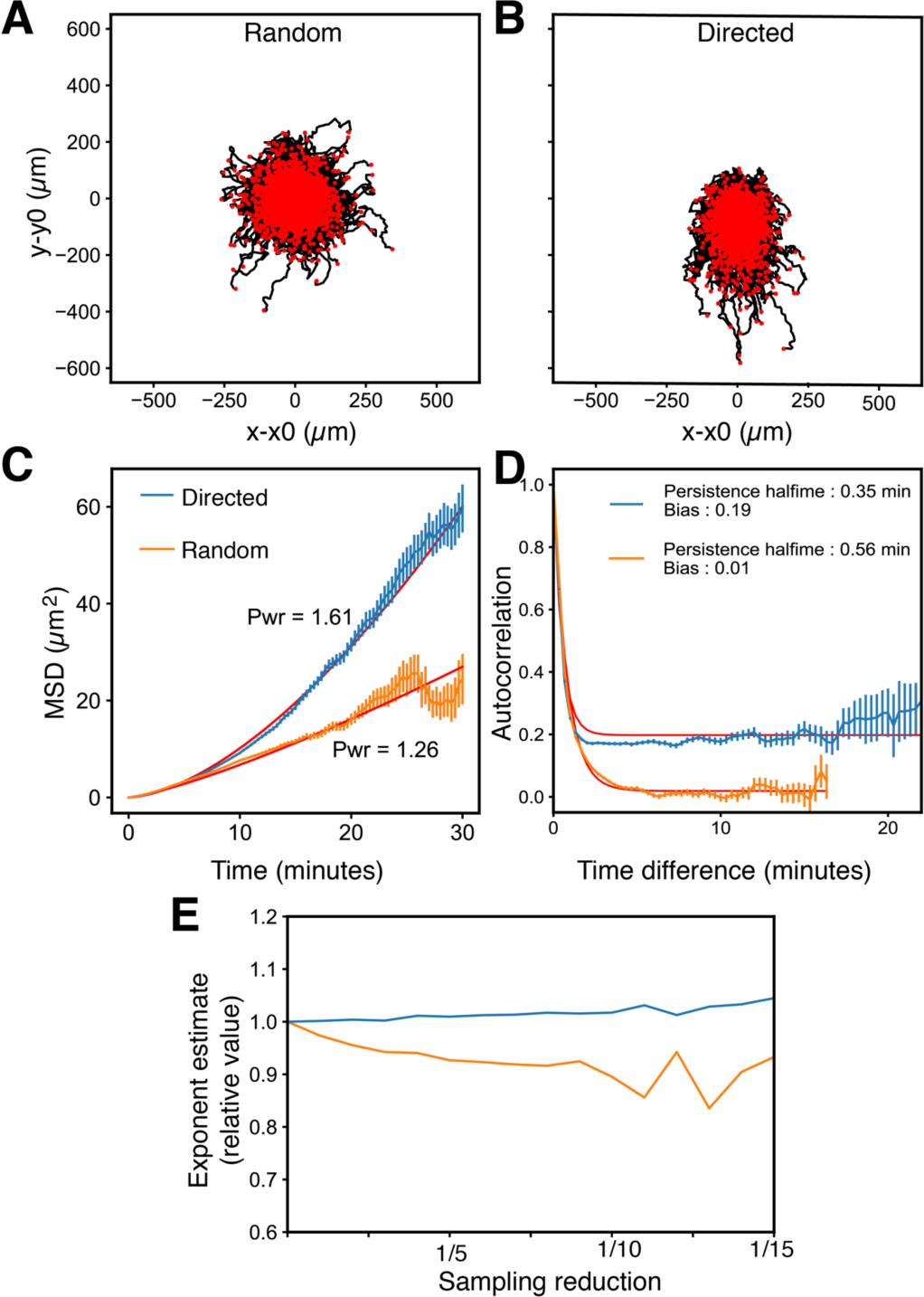
MSD and autocorrelations of real chemotaxing cells. **(A, B)** Spider plots of cells migrating to a uniform (A, ‘Random’) or a directional (B, ‘Directed’) attractant stimulus under agarose. **(C)**, MSDs for the random (orange) and directed (blue) assays in real cells. The directed migration assay is highly ballistic. The random migration assay, though it has a much low exponent, remains superdiffusive, meaning that cell behaviour does not correspond well to a theoretical random walk. **(D)** Autocorrelations for the random (orange) and directed (blue) assays. **(E)** The values of MSDs for the random (orange) and directed (blue). As frames are dropped from the analysis. The values remain very consistent even when far fewer frames are available, demonstrating a robustness in this method to the sampling rate.

### Transition matrices capture complex persistence and bias

A third informative way to distinguish persistence and bias is provided by creating a transition matrix (Taylor et al., 2013) examining the relationship between direction of motion at a given time with the direction of motion a short time later. In these plots the Y axis shows the direction that a cell is moving in, and the X axis shows its direction in the previous frame (Fig 4A). In simple, randomly moving cells there is no correlation between their direction and the stimulus, so points are spread evenly over the plot (Fig. 4B, TL). Biased paths tend to cluster at a single point (in this case the centre, 0°), as cells’ directions of motion cluster around the direction of the stimulus (Fig. 4B, TR). Similarly, when cells move persistently, their paths are represented in the diagonal from the bottom left to the top right; no one direction dominates but the direction of motion changes very little between frames. A diagonal stripe therefore typifies the PRW (Fig. 4B, BL). The PBRW has the features of both, with a bias hotspot skewed along the diagonal by directional persistence (Fig. 4B, BR). These features can be readily seen in real data: the transition matrices for the non-directional (Fig. 4C, top) and directional (Fig 4C, middle) show clear similarities with the persistent- and persistent biased random walks, respectively, though the latter displays a hotspot in a different position as the cells are migrating downwards(-90°) in the experiment, rather than to the right (0°). Finally, transition matrices can record more complex chemotactic behaviours than either summary statistics or model fitting-Fig 4C, bottom, shows cells stimulated by an initially uniform cAMP stimulus in a chamber. As they degrade the attractant in the assays, the cells are attracted outward, with each cell moving to the closer of the two attractant reservoirs. This creates two hotspots on the corresponding transition matrix, one for the cells heading upwards, another for the cells heading downwards.

**Fig. 4:**
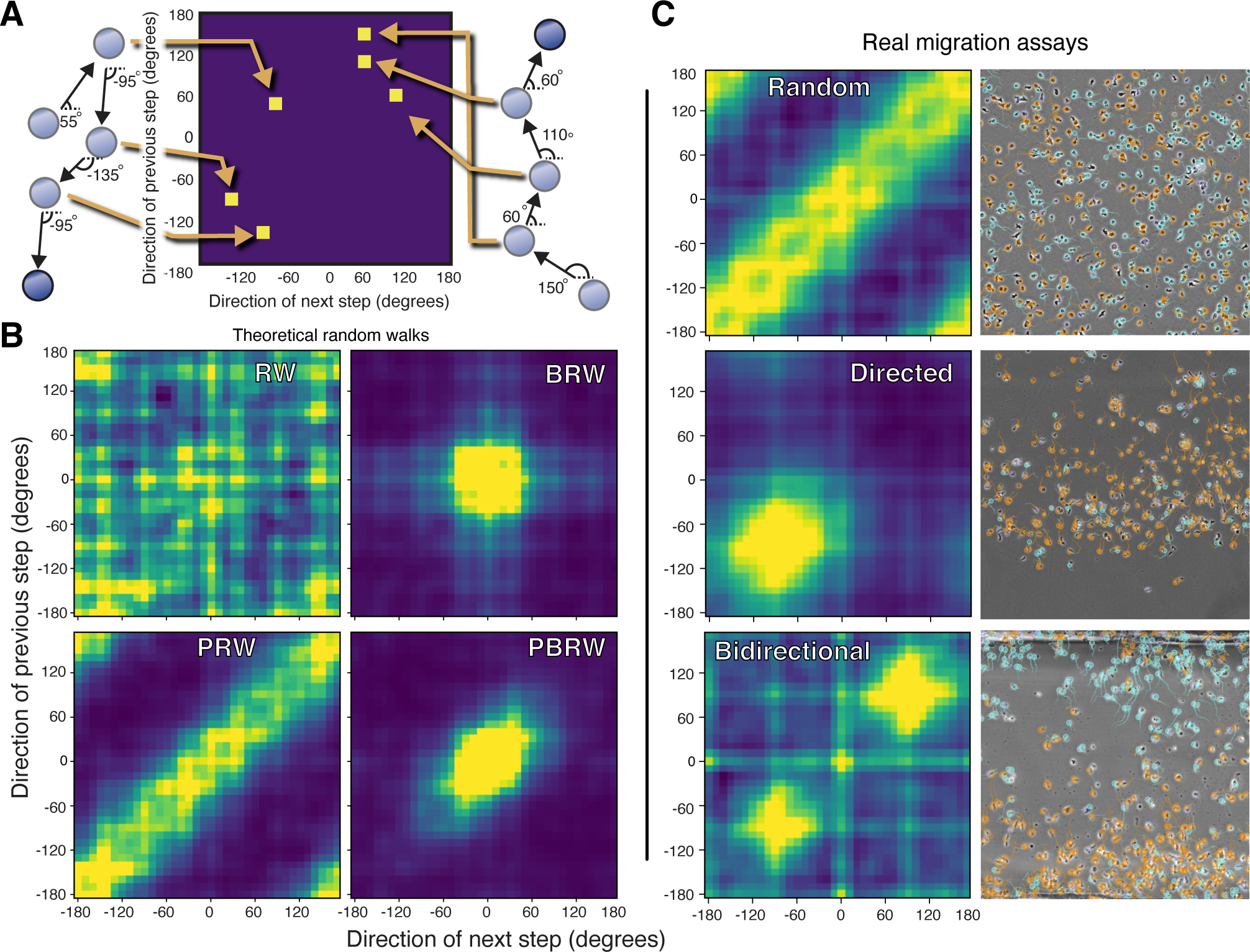
Transition matrices help visualise underlying behaviour. **(A)** Schematic example for making a transition matrix. At each point in a cell track except the first and last, the absolute angle to the next point on the track provides the x-axis coordinate and the absolute angle to the previous point provides the y-axis coordinate. A heat map is built up of all the recorded movements to show which directions are favoured given the previous direction of movement. **(B)** Transition matrices for the four theoretical random walks. **RW:** There is no relationship between the direction at one time and another. This makes the transition matrix quite homogeneous, with bright areas unclustered. The gridding effect is an artefact of the discrete co-ordinate system in the image in which the cells were tracked (i.e. cell position is measure to the pixel). **BRW:** A clear bright spot dominates at a particular point along the upward diagonal. This angle will be the same on both axes, and will be the direction of bias in the image (in this case, 0 degrees). **PRW:** As cells persist in a direction between frames, the transition contains a bright upward diagonal band, showing little change in direction. If this is absent, it does not mean that there is no persistence but that the persistence time is shorter than the sampling interval. **(PBRW)** The persistent biased random walk looks like a biased random walk, but with the directional hotspot stretched along the “persistent” upward diagonal. **(C)** Transition matrices for real examples. **Random:** Uniformly distributed *D. discoideum*, starved and stimulated with Sp-cAMPS, producing a PRW effect. Directed: The same cells in a one-well under agarose assay to cAMP. Their movement is directed at -90° and appears to be a PBRW. **Bidirectional:** The same cells, uniformly distributed between two attractant-rich reservoirs in an Insall chamber. As the cells degrade this attractant, they create a gradient leading away from the centre and split down the middle, migrating toward their nearer reservoir.

One further advantage of transition matrices is that they display a distribution of observations concisely, making the quality of the data apparent. The disadvantage is that they do not yield quantitative parameter measurements. They are therefore best used together with MSD and/or autocorrelation.

### Measurement artifacts caused by nonhomogeneous distribution of cells

Fitted models work well when dealing with full information or an unbiased ensemble of quantitated paths. They are not robust to some forms of bias that are common in real experiments (this is true for many types of measurement). For example, in many chemotaxis chambers (Zigmond, Dunn, Insall, Ibidi) the standard way of setting up the assay involves cells attached homogeneously over the bridge. This makes it possible to observe cells moving in all possible directions, so the datasets are unbiased. However, in transwell assays the cells are introduced to one side of the chamber, and their measurement to the other side is counted; only one direction of migration can be observed. Direct-view chambers can be used in a similar way, particularly when observing self-generated chemotaxis (Tweedy et al., 2016), with cells introduced through one well, and none initially on the bridge. Cells are only observed as they move out from the well to the bridge, and those with high persistence will continue in this direction regardless of bias. This can give an appearance of chemotaxis, because calls that move randomly in other directions will not arrive on the bridge and not be observed. Fig. 5A shows the automatic tracking of an unbiased, persistent walk starting in a staging area for the whole domain (top), as well as for two smaller regions to the right of the staging area. Though the true transition matrix is that of a persistent random walk (Fig. 5B, top panel), our smaller observation region does not include those cells that initially move toward the left, and so the transition matrix looks like that of a persistent biased random walk (Fig. 5B, middle and lower panel). More traditional analyses such as Spider plots and Cos(8) show the same bias (Fig. 5C), and even directional autocorrelations register a false positive (Fig. 5D). We must show care when interpreting the results of experiments that could introduce these biases (e.g. always including a paired experimental negative control, rather than comparing behaviour against a statistical model of neutral behaviour).

**Fig. 5:**
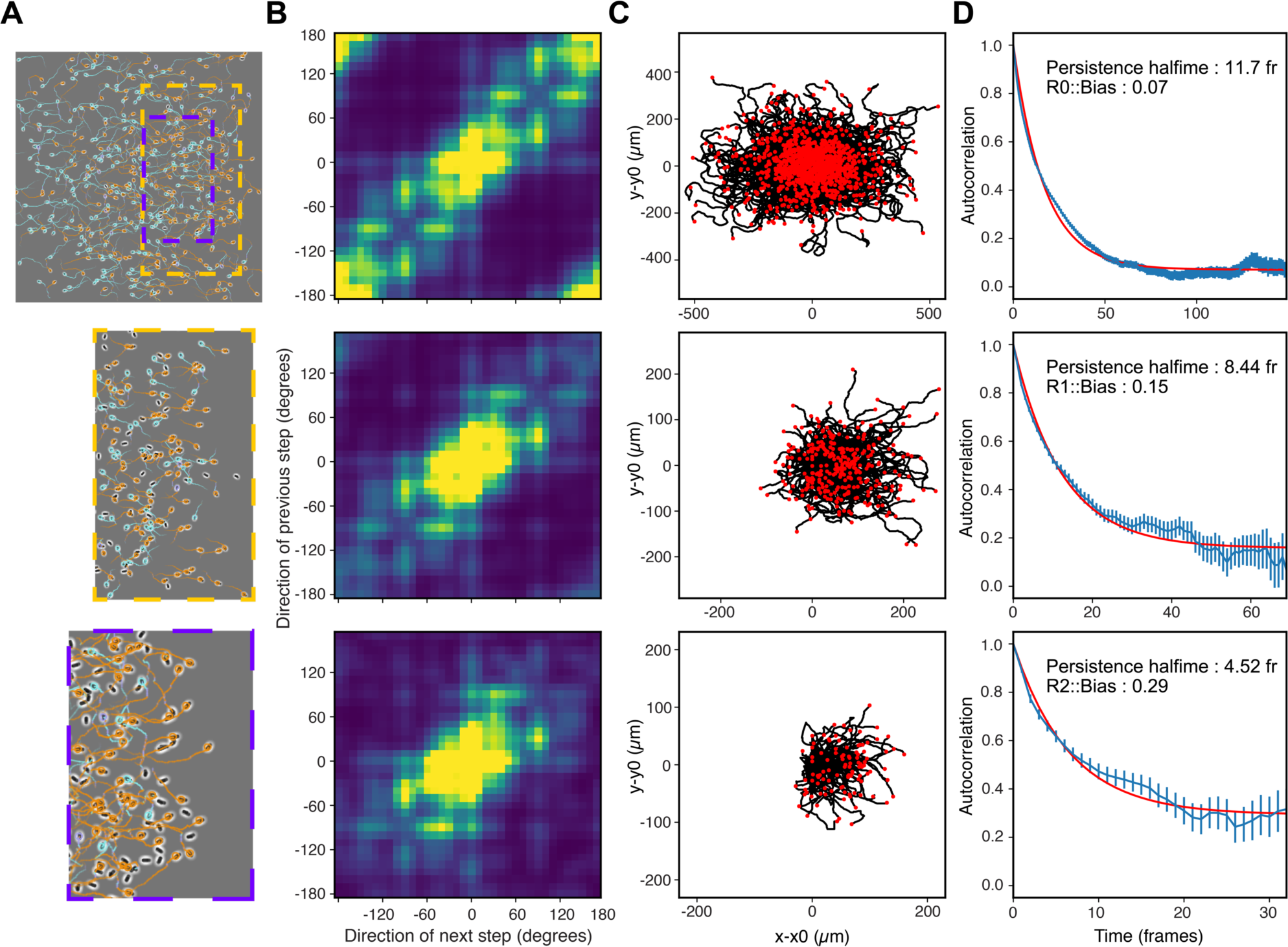
Non-homogeneous initial conditions can bias statistics. **(A)** A random walk with high persistence and no bias. We compare the results without any bias in our field of view (top), with two situations in which we can only see the area immediately to one side of where the cells start (shown on the original image with orange and purple dashed boxes). **(B)** The transition matrices for each example. Note that those with a biased field incorrectly represent the cell movement as biased. **(C)** Spider plots and Cos(**8**) values for each case. **(D)** Autocorrelations. Surprisingly, even the fitted bias parameter here is incorrectly amplified.

### Real biology breaks assumptions: receptor saturation, attractant alteration and individual vs ensemble experience

Eukaryotic cell migration is complicated and often counterintuitive. This means that simplifying assumptions can be incorrect, and experimenters must be wary of spurious conclusions. Cells that are chemotaxing under physiological conditions may also be influenced by chemokinesis (Zigmond, 1977), attractant degradation by cells as they migrate (Tweedy and Insall, 2020), and secondary signals secreted by cells responding to attractants (Devreotes et al., 1979, Lammermann et al., 2013, Zaki et al., 2006). In addition, when cells are observed chemotaxing over time, they experience changes in the concentrations of attractants and receptor saturation as they climb the gradient. Even in the most relied-on assays, we notice features that make experimental data deviate from both the straightforward descriptions of the outcomes, and the simplified theories in Fig. 2 most commonly used to describe them.

For example, during migration of *D. discoideum* to an imposed gradient of cAMP, cells are initially spread uniformly between a reservoir of attractant to the right, and a sink containing only medium to the left, and expect a chemotactic response to a linear gradient that diffusively forms between these two features (co-diffusing fluorescent markers have sometimes been used alongside the attractant as a gradient visualisation tool; Veltman et al., 2008). We repeated this experiment for the purposes of this article, and measured movement of cells over the entire chamber. The directional movement of cells in the original movie is clear (Video 1), but Cos(8) is poor and autocorrelation reports next to no bias (Fig 6A, B, upper panel). Equally clear is that directional migration is restricted to the right-hand side of the viewing area, closest to the attractant reservoir (Fig. 6C, upper panel). Cells further down show no signs of directionality. When our model includes this attractant degradation, the observed behaviour makes sense. Cells closest to the attractant reservoir see a steep gradient, but little or no attractant diffuses through to the cells further away. This illustrates that the experiment as envisioned can differ strongly from the experiment as it functions. The complexity that this cell-environment interaction adds is missing from the canonical random walks. A cell moving across the whole domain will behave differently at different times through its migration, and its track will not fit to any fixed model. In addition, if we include the cAMP degrading ability of real cells in our model of this assay (conceptually or computationally), we will predict an attractant gradient that is approximately exponential, rather than linear (Tweedy et al., 2016).

**Fig. 6:**
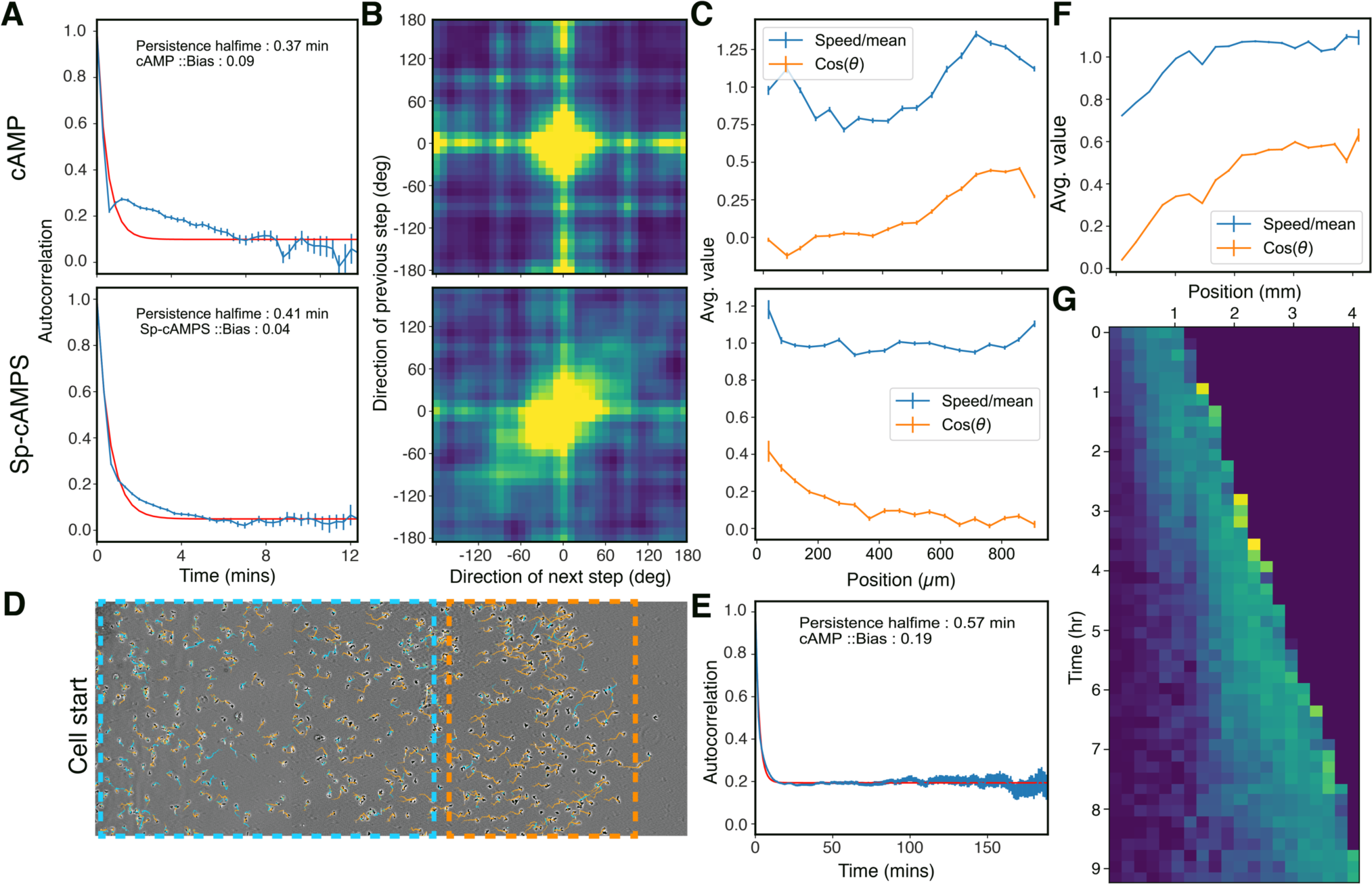
Real experiments disagree with simple model behaviour. **(A)** Experiments performed with *D. discoideum* toward gradients of 10µM cAMP and 1µM Sp-cAMPS, imposed in a chemotaxis chamber across 1mm. Although this is a standard, well optimised chemotaxis experiment, the measured bias of the track ensemble is low in both cases. **(B)** Transition matrices for both gradients. **(C)** Taking averages of Cos(**8**) in 50µm bins across the 1mm bridge, we see that neither behaves uniformly. In the cAMP case, Cos(**8**) is high at the top of the bridge and close to zero across the bottom half. The attractant diffusing in from the top end of the gradient is degraded before reaching the cells in the bottom half. This is a powerful demonstration that cells can shape their environment, and that the gradient imposed by the experimentalist is not necessarily that which is seen by the cells. In the Sp-cAMPS case, the opposite profile is seen, with the highest values of Cos(**8**) at the low end of the bridge. As Sp-cAMPS is not degraded, it is reasonable to assume that the cells are responding to the initially imposed gradient. The deviations here are caused by receptor saturation at the high end. This can only be avoided by lowering the overall gradient steepness, reducing chemotaxis at the low end. **(D)** Self-generated gradient assay to an initially uniform 10µM folate background. Tracks for the previous 10 frames are shown, with positive directionality shown in orange and negative shown in light blue. **(E)** Autocorrelation of the assay in (D). **(F)** Positional binning of the assay shown in (D) shows the same pattern as the cAMP (C). This makes sense as, in both cases, the attractant is degraded, however this analysis does not properly describe the behaviour of the cell ensemble. **(G)** Heat map of projected speed, binning position (y axis) and time (x axis) shows a front of cells advancing more or less linearly over time. Also noteworthy is that, behind this front of cells, directionality is lost (purple area bottom & right). These two behaviours can be seen in (D), with the front outlined in an orange box, and the rear outlined in a light blue box.

To discount the effects of chemoattractant breakdown, we set up the same experiment, but in place of cAMP we use a non-degradable (Theibert et al., 1986) analogue, Sp-cAMPS (Fig. 6A, B, lower panel; Video 2). Again, taking an average of all cells, we see poor chemotaxis, but this time when we look at the profile of behaviour across the bridge we see the reverse of our cAMP results: good chemotaxis is confined to the area furthest from the attractant well (Fig. 6C, lower panel). As Sp-cAMPS is not degraded, the absolute gradient is linear, however near the attractant well the concentration is higher. As the receptors become more saturated across the cell, the number available for an additional response at the front becomes lower, so the differential between the numbers of receptors activated at the front and back drops. At high receptor occupancies, the response is saturated, and no difference can be distinguished between the front and the rear. This means that cells at the low end of the gradient can perceive a cAMP gradient much better than those at the high end (Dowdell et al., 2023). This phenomenon has led to SpCAMPS being called a weak attractant (Van Haastert et al., 1982), in spite of evidence that it is a near-perfect agonist for the cAMP receptor (Van Haastert et al., 1984). In both of these cases, a thorough spatial analysis (rather than simply reporting a migration statistic) is crucial to understanding the biology and the limits of our current models of chemotaxis.

A special case of attractant degradation is the self-generated gradient, in which cells reshape a uniform attractant profile into a steep, directional gradient cue (Insall, 2023). One key feature of self-generated chemotaxis is the emergence of a sharp, self-sustaining wave of chemotaxis, accompanied by a steep local attractant gradient. The cells ahead of the wave chemotax weakly because of receptor saturation, and cells behind the wave do not chemotax because all the attractant has been depleted. The cells within the wave, however, chemotax exaggeratedly well to the locally steep gradient; and the behaviour of the entire population is completely shaped by chemotaxis, even if counterintuitively many of these cells are undirected at any one time of observation (Tweedy et al., 2016). Fig 6D shows automatic tracking for one example, *D. discoideum* migrating to folate. The assay is similar to that shown in Fig. 4C (directed), but imaged over a greater distance. Autocorrelation and Cos(8) show a moderate bias overall (Fig. 6E), but these statistics miss the most interesting features of cell behaviour in this assay, which are extremely localised, as does binning the data across space as in Fig. 6A. In the latter case, time-averaging the profile causes us to miss the dynamic behaviour in this system. It merges the behaviour of those cells that arrived first, chemotaxing past and degrading attractant, with those that were left behind, which randomly migrate as there is no longer any attractant to respond to.

In this assay, correct analysis of cell behaviour in space and time is best performed using a heatmap, binning cells’ projected speed in both dimensions (Fig. 6G). This heatmap shows three regions-the area as yet unreached by cells (Fig. 6G, top left purple area), the wave of cells chemotaxing forward (bright diagonal, bottom left to top right), and finally the cells left behind which move only randomly (purple noisy area, bottom right). In reporting only Cos(8), or autocorrelation bias, practically all of the interesting information would be lost, reduced to a single number that tells us nothing more than “cells go rightwards”, and implying general weak chemotaxis instead of the locally specific excellent steering.

## Discussion

Single value statistics will always have an important role in studies of cell migration. They are easy to calculate and allow for the straightforward comparison of different experimental conditions. Nonetheless, many recent advances in understanding cell migration can be actively hidden when complex data are reduced to a single parameter. These are advances defined by a complexity that is lost by reporting only gross trends, and their complicating factors are what makes them valuable.

Model fitting can be more robust, and well-fitted parameters come with easily interpreted meanings that paint a richer picture than any single population statistic. Obviously, model fitting requires rich enough data, meaning that direct view chambers rather than e.g. transwell assays are required, and such approaches should not be assumed to work off-the-shelf. Different cell types exhibit a startling range of complex behaviours that can frustrate understanding via model fitting, unless the right model is chosen. Secreting, degrading and modifying attractants, chemokinesis and contact inhibition are all examples of common cell behaviours, and no two are mutually exclusive. In our extensive work on cell migration, we know of no case that fits to the simple model of chemotaxis entirely. There are many more models than the one discussed here, and each experimental regime has appropriate ones; in particular, Levy walks are often used in modelling cell and animal migration. However, wider use of the tests we describe in this paper would greatly improve the interpretation of most data and our understanding of chemotaxis.

When we design an experiment so that it is practical to perform, we must account for the introduction of biases. These could be mutations that make cells easier to cultivate under lab conditions, for example, or microscopy choices that make imaging easier. Here, we showed that biases can be introduced by the choice of observation position relative to the centre of a population of cells. Even if these biases are subtle, some statistical measurements, such as Rayleigh tests, are highly sensitive and may be prone to false positives introduced by experimental technique. This demonstrates the necessity of a full suite of experimental controls. It is important to be thoughtful about what this means, and consider controls other than a negative – for example, heightened chemokinesis relative to a negative control could easily be misidentified as chemotaxis in an inhomogeneous field of cells.

Finally, while there will continue to be a place for single-valued population statistics under appropriate circumstances, our analysis shows that Cos(8) probably ought never to be used. It is ambiguous, and encourages ambiguity. The better version of Cos(8), in which multiple steps are averaged, is extremely dependent on the frame rate and the relative cell speed. Overall, it has no advantages over the chemotaxis efficiency index, CEI, which is as simple to calculate and far more robust. We urge members of the field to use CEI, rather than ever using Cos(8), and have provided an ImageJ plugin that comes with this paper to help its uptake.

## Methods

### Cells & Reagents

Chemicals were obtained from Sigma-Aldrich except where stated. The cells used in this work were wild Dictyostelium discoideum of the strain NC4. Cells were cultured on bacterial lawns on SM agar (from Formedium, https://formedium.com/). For assays using cAMP/Sp-cAMPS (Fig 3, 4, 6A&B; Sp-cAMPS from BioLog Life Sciences Institute, https://www.biolog.de/), cells were developed on development buffer (DB) agar until scrolling waves were observed by visual inspection, and were then collected, pelleted, and resuspended in experimental medium (DB+3mM caffeine). Experiments used caffeine to supress cAMP release by the cells. For experiments using folate (Fig. 6D), cells were taken directly from growth plates, washed (3×300g, 3min) and resuspended in experimental media (10mM folate in LoFlo medium, Obtained from Formedium, https://formedium.com/).

### Chemotaxis experiments

Experiments were in Insall chambers or under agarose. Insall chambers were set up in one of two ways: 1) with cells settled on a glass coverslip, then inverted onto the chamber and imaged in the viewing area where the gradient forms [Fig. 4C, Bidirectional, Fig. 6A-C]. 2) With cells added to the outside well and imaged as they migrate in toward the centre well (Fig. 2B, 3C Directed). In the random migration condition (Fig. 3A), developed cells are settled in the wells of a 6 well dish and placed under a slab of agarose containing a non-degradable attractant (20mM Sp-cAMPS). In the self-generated gradient assay (Fig. 6D), vegetative cells are placed in a single well cut into LoFLo agarose containing the attractant (10mM folate). All cells were imaged used a Nikon Ti2E microscope, phase contrast using a 10x 0.4NA objective and Retiga R6 camera, with typically one frame every 30 seconds.

### Simulations

As previously described [MAZES?]. Briefly-cells move continuously against a discrete background grid, which simulates attractant diffusion using the Dufort Frankel method. Cell movement is a persistent biased random walk, with the bias supplied by the slope of receptor occupancy in the local region of the cell. Receptors are occupied at their dynamic steady state and saturate with simple, first order kinetics. Where cells degrade attractant, they do so with Michaelis Menten kinetics.

### Tracking and analysis

These aspects were chiefly performed using a custom-written plugin for ImageJ. Points are found using the in-built ImageJ local maximum finder and tracking uses constrained point-matching algorithm with an L2 cost. The plugin is written in Java by LT. Source code is available on Github at [to be added upon publication]. Analysis is also performed in python, with the figures in this paper produced using matplotlib.pyplot. The relevant code is also available via Github [to be added upon **publication].**

## Acknowledgements

We are very grateful to Dr. Peggy Paschke for substantial advice on the required features of our ImageJ tracker, Prof. Laura Machesky for comments on the manuscript, and to the Beatson Advanced Imaging Resource (BAIR) facility for help with microscopy.

## Author contributions

**LT**: Conceptualisation, Experiments, Investigation, Software, Analysis, Visualization, Writing; **PT**: Conceptualisation and editing; **RHI**: Conceptualization, Funding acquisition, Supervision, Writing.

## Video Legends

**Video 1: Cells chemotaxing to degradeable attractant in a direct-view chamber** Developed *Dictyostelium* cells were exposed to a gradient of 0-1µM cAMP in an Insall chamber, imaged through a 10x Nikon objective and filmed using a Retiga R6 camera. The higher concentration is shown to the right. cAMP is rapidly degraded by a secreted phosphodiesterase. Frame rate 1/15s.

**Video 2: Cells chemotaxing to non-degradeable attractant in a direct-view chamber** Developed *Dictyostelium* cells were exposed to a gradient of 0-10µM Sp-cAMPS in an Insall chamber, imaged through a 10x Nikon objective and filmed using a Retiga R6 camera. The higher concentration is shown to the right. Sp-cAMPS is a cAMP analogue that induces chemotaxis but cannot be degraded by phosphodiesterases. Frame rate 1/15s.

